# Changes in nanomechanical properties of single neuroblastoma cells as a model for oxygen and glucose deprivation (OGD)

**DOI:** 10.1101/2022.02.19.481155

**Authors:** Tomasz Zieliński, Joanna Pabijan, Bartłomiej Zapotoczny, Joanna Zemła, Julita Wesołowska, Joanna Pera, Małgorzata Lekka

## Abstract

The biological processes underlying ischemic stroke, although complex, are better known than those related to biomechanical alterations of single cells. Mechanisms of biomechanical changes and their relations to the molecular processes are crucial for understanding the function and dysfunction of the brain. In our study, we applied atomic force microscopy (AFM) to quantify the alterations in biomechanical properties in neuroblastoma SH-SY5Y cells subjected to oxygen and glucose deprivation (OGD) and reoxygenation (RO). Obtained results reveal several characteristics. Cell viability remained at the same level, regardless of the OGD and RO conditions, but, in parallel, the metabolic activity of cells decreased with OGD duration. 24h RO did not recover the metabolic activity fully. Cells subjected to OGD appeared softer than control cells. Cell softening was strongly present in cells after 1h of OGD and, with longer OGD duration and in RO conditions, cells recovered their mechanical properties. Changes in the nanomechanical properties of cells were attributed to the remodelling of actin filaments, which was related to cofilin-based regulation and impaired metabolic activity of cells. The presented study shows the importance of nanomechanics in research on ischemic-related pathological processes such as stroke.

## Introduction

Ischemic stroke remains one of the leading causes of death, especially in the elderly^1^. It is caused by disrupted blood flow to the brain resulting in oxygen and glucose deficiencies in the cells. The last three decades show significant improvements in acute treatment, resulting in increased life expectancy after treatment and rehabilitation^1, 2^. Understanding the stroke at the cellular level can be simulated using an in vitro oxygen-glucose deprivation (OGD) model. The model was widely investigated to study ischemic cell death^3^. In the model, cells or tissue slices are exposed to hypoxic or anoxic conditions and cultured in media deprived of glucose. Not only the effect of OGD is investigated in the model – after changing media and introducing normal oxygen levels, reperfusion can be additionally tested. With long-lasting OGD, the reoxygenation may paradoxically cause additional damage. Ischemia-reperfusion injury is caused by the immediate generation of reactive oxygen species, altered ion transport, and calcium influx^4^. During OGD, rapid remodeling of the actin cytoskeleton was reported to be involved in the blood-brain barrier disruption and affected: endothelial cells^5^, non-neuronal brain cells^6^, and neurons^7^.

Actin filaments occur in a cell as a meshwork or bundles of parallel fibers abundant, particularly close beneath the cell membrane^8, 9^. The continuous control of the balance between polymerization and depolymerization ensures a dynamic equilibrium state, controlling cell architecture, mechanical resistance, and regulating many biological processes^10^. The dynamic of this process is regulated by actin-binding proteins^11^. Cofilin, an actin-depolymerizing factor, was highlighted several times to play a crucial role in actin remodeling in axons^7, 12, 13^. In ischemia-induced actin disruption, cofilin was linked with ATP depletion^14^. It has already been reported that cofilin is essential for an early phase of apoptosis^15^ or intracellular contractile force generation^16^. The responsibility of cofilin and its role in various diseases makes it a potential target for potential neuroprotective approaches in the early stages of ischemic brain injury. In particular, the SH-SY5Y human neuroblastoma cell line is used to investigate the OGD model of stroke^17^. The cell line is of human origin, allowing for a better reflection of the induced changes during the stroke. Both non-differentiated and differentiated SH-SY5Y cells have their advantages and drawbacks in the model of neuron cells^18^. In this report, we used undifferentiated cells, which are considered to be most reminiscent of immature neurons^19, 20^.

In the present study, we hypothesize that possible involvement of cofilin occurs in the early stages of cytoskeleton remodelling under ischemic conditions. In the initial phase, such remodelling is limited to actin filaments reorganization, which can be quantitatively evaluated using an atomic force microscope (AFM)^21^. This technique is characterized by nanoscale resolution enabled to quantify fine alterations in cells and tissue nanomechanical properties in normal and pathological conditons^22–25^. The changes in mechanical properties have already been shown in undifferentiated SH-SY5Y cells in a model of chemically induced neurodegeneration^26^. The results, associated with glutamate-mediated neurodegeneration, showed the increased rigidity of SH-SY5Y cells upon 50 mM N-methyl-D-aspartate (NMDA) treatment. Although the experiment time was limited to 60 min, the maximum rigidity values were obtained after 20 minutes^26^. Thus, NMDA induced cytoskeletal reorganization. However still, knowledge about the mechanical changes in OGD/reoxygenation (RO) is lacking. Thus, we analyzed the nanomechanical properties of SH-SY5Y neuroblastoma cells exposed to oxygen and glucose deprivation, mimicking ischemic conditions. Following our previous studies on the effect of anti-tumor drugs on prostate cancer cells^27^, two indentations were applied, i.e., shallow (400 nm) and deep (1200 nm) ones. The shallow indentation reveals mechanical properties dominated by actin filaments, while the deep indentation may contain the additional contribution from deeper parts of the cells like the microtubular network and cell nucleus^27, 28^. The studies were accompanied by evaluating the cofilin and phosphorylated cofilin expression levels, visualization of actin filaments organization quantified using morphometric parameters, and metabolic activity of SH-SY5Y cells subjected to OGD. Measurements were conducted directly after OGD to study the magnitude of the induced changes and after 24h of reoxygenation to model reperfusion and to evaluate the reversibility of these changes.

## Results

### Viability of SH-SY5Y cells under OGD

To assess the effect of OGD exposure (5% CO_2_, 0.1% O_2_) on neuroblastoma SH-SY5Y cells, we exposed SH-SY5Y human neuroblastoma cells to OGD for 1, 3, and 12 hours, followed by 24 hour-RO. (**Fig. 1**).

**Figure 1.**
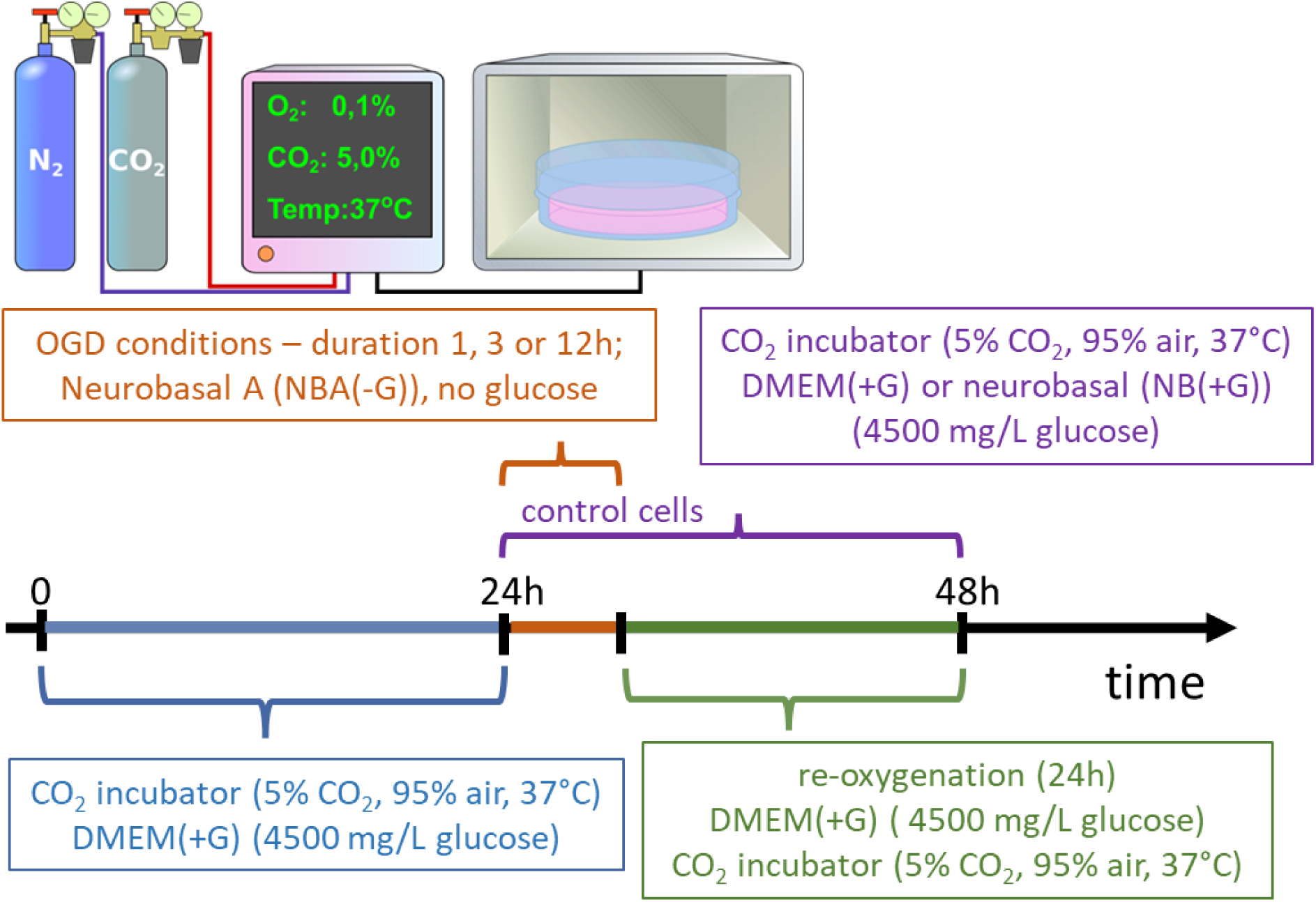
(a) A scheme showing three steps of sequential OGD applied to living SH-SY5Y cells. Firstly, cells were cultured for 24 hours after seeding in 5% CO_2_, 95% atmosphere (37°C) in a DMEM with 4500 mg/ml of glucose (DMEM(+G)). They refer here as control cells. Next, the medium was exchanged to NBA(-G), and cells were placed in a table CO_2_ incubator for 1h, 3h, or 12h at 0.1% O_2_ (referred to as OGD conditions and OGD cells). Finally, OGD cells were rinsed with a DMEM(+G) in the atmosphere of 5% CO_2_ and 95% air (reoxygenation conditions, , in addition, non-OGD cells were kept in DMEM(+G)). **(b)** Phase-contrast image showing the morphology of neuroblastoma SH-SY5Y cells cultured for 24h in NB(+G), as it induced differentiation resulting in a neuron-like morphology with numerous, fine protrusions (neurite-like structures). Scale bar – 50 µm.

We compare four groups of data, namely, control (C, measurements were conducted in neurobasal medium, which contained 4500 mg/L of glucose, referred here as NB(+G)), OGD cells (in neurobasal A medium without glucose, NBA(-G)), reoxygenated OGD cells (DMEM, which contained 4500 mg/L of glucose, DMEM(+G)) and additional control (i.e. non-OGD) cells kept in DMEM(+G) for the same time as re-oxygenated (RO) OGD cells.

We started with the assessments of metabolic activity (using MTS assay; reduction of tetrazolium; impaired NAD(P)H metabolism^29^) and cell viability (using LDH assay; lactate dehydrogenase release to culture media, membrane damage^30^) that were applied to samples collected directly after OGD and after 24h of reoxygenation. The results show that cell metabolism and viability depended on OGD duration (**Fig. 2**). Moreover, the induced changes are still present in cells after reoxygenation.

**Figure 2.**
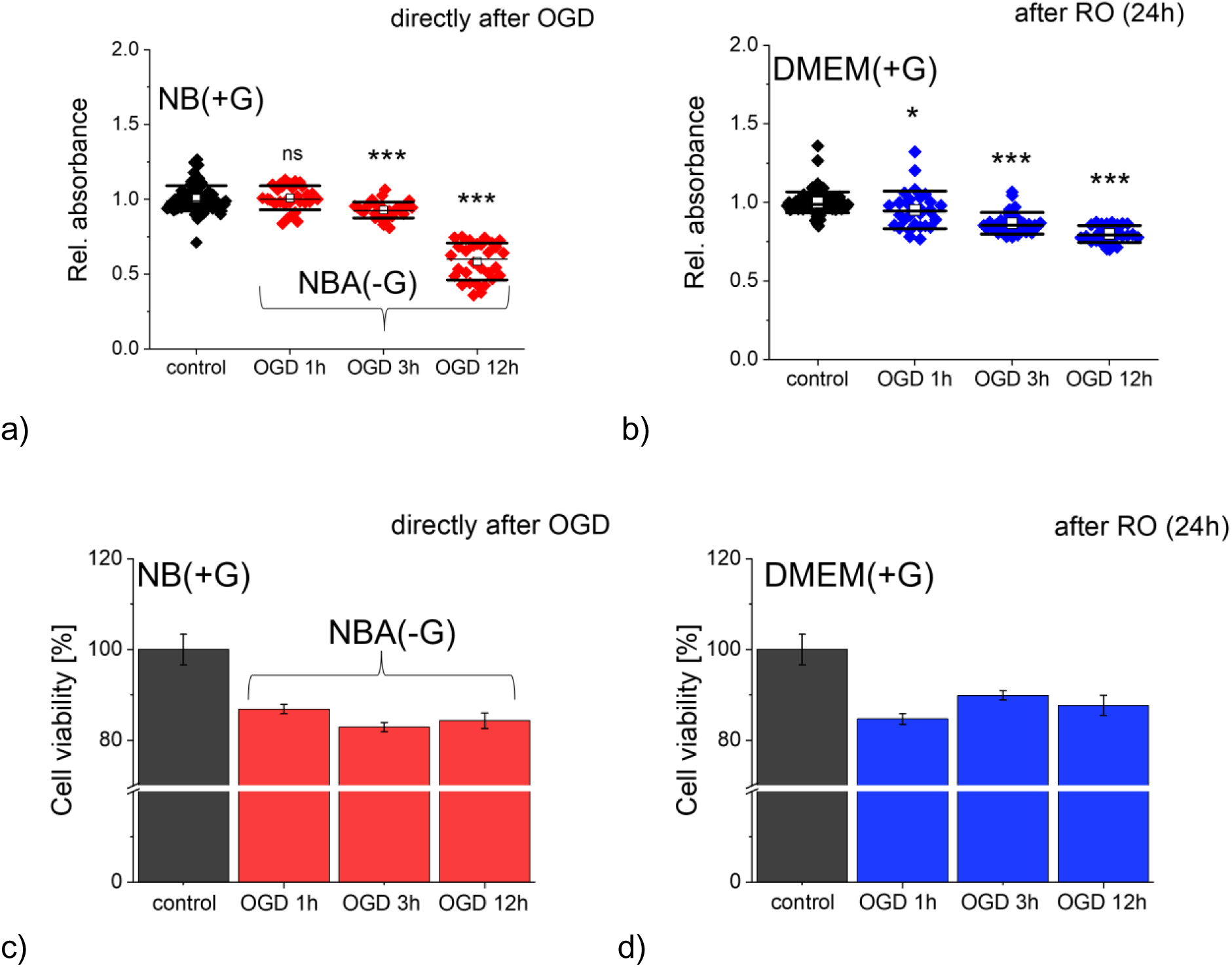
Metabolic level and viability of SH-SY5Y neuroblastoma cells assessed by MTS (a,b) and LDH (c,d) assays, directly after OGD (a,c) and after 24h of reoxygenation (b,d). Each dot denotes a single readout from the ELISA reader. (a, b) A mean (open circle), median (black line), standard deviation (SD, box size) were determined from data gathered from 3 independent repetitions. (c,d) Columns represent a mean value from 12 ELISA readouts (*n* = 3 independent repetitions). Relative absorbance was normalized to values obtained for the control samples. Statistical significance: *ns* – not statistically significant, *p* > 0.05, *\*p* < 0.05, *\*\*\*p* < 0.001.

We tested how 1h, 3h, and 12h of OGD affect cell metabolism of SH-SY5Y cells (**Fig. 2ab**). The MTS tetrazolium is reduced by cells to formazan soluble in the culture medium. Such conversion is presumably accomplished by NADPH or NADH produced by dehydrogenase enzymes in metabolically active cells^29, 31^. Thus, lower absorbance in comparison with control cells denotes the lower metabolic activity of cells. Our results show a significant reduction in formazan conversion after 3h and 12h. After one hour of the cell exposure to OGD, the metabolic activity level was similar to that of control cells, and no significant difference was identified (*p* = 0.262; **Fig. 2a**). However, we do observe changes in reoxygenation for all three groups of cells subjected to OGD. The MTS-based cell viability assessed directly after OGD experiments dropped by about 7.1% (*p* < 0.001) and 41.5% (*p* < 0.001) after 3h and 12h of cell exposure to OGD, respectively (**Fig. 2b**). In parallel, we checked membrane integrity by LDH assay related to the number of viable cells. A drop of about 13% – 17% was observed for cells after OGD (**Fig. 2c**). A similar drop in the number of viable cells was observed for cells kept for 24h in RO. A drop between 12%-15% was obtained (**Fig. 2d**).

We expect that cells being damaged by OGD will recover their ability to proliferate^32^. Therefore, both MTS and LdH assays have also been applied to cells cultured in reoxygenation conditions. MTS results revealed significant changes in all groups of cells (**Fig. 2b,d**). The most significant drop in cell metabolic activity was observed for cells subjected to OGD for 12h, while a small change, but still statistically significant, was recorded for cells after 1h of OGD. Interestingly, the level of LdH remained at a similar level for both control and treated (OGD, RO) cells (**Fig. 2c,d**), indicating no correlation between cell viability and metabolic activity of cells, regardless of their treatment.

These results show that only metabolic activity was affected after prolonged OGD. Lower metabolic activity was not related to the number of viable cells (or more precisely to the impaired membrane integrity). No new cells were dying after 24h OGD, but their metabolism activity was altered after OGD.

### The effect of different OGD duration on mechanical properties of SH-SY5Y cells

To assess whether the altered metabolic activity is related to nanomechanical properties of SHSY%Y cells, AFM working in a force spectroscopy mode was employed to conduct the measurements over a nuclear region of the cell (to avoid the influence of stiff substrates^33^). The nanomechanical properties were quantified by Young’s (elastic) modulus calculated by applying Hertz-Sneddon contact mechanics^22, 34^, assuming that a cone can approximate the shape of the probing pyramidal tip (**Fig. 3**).

**Figure 3.**
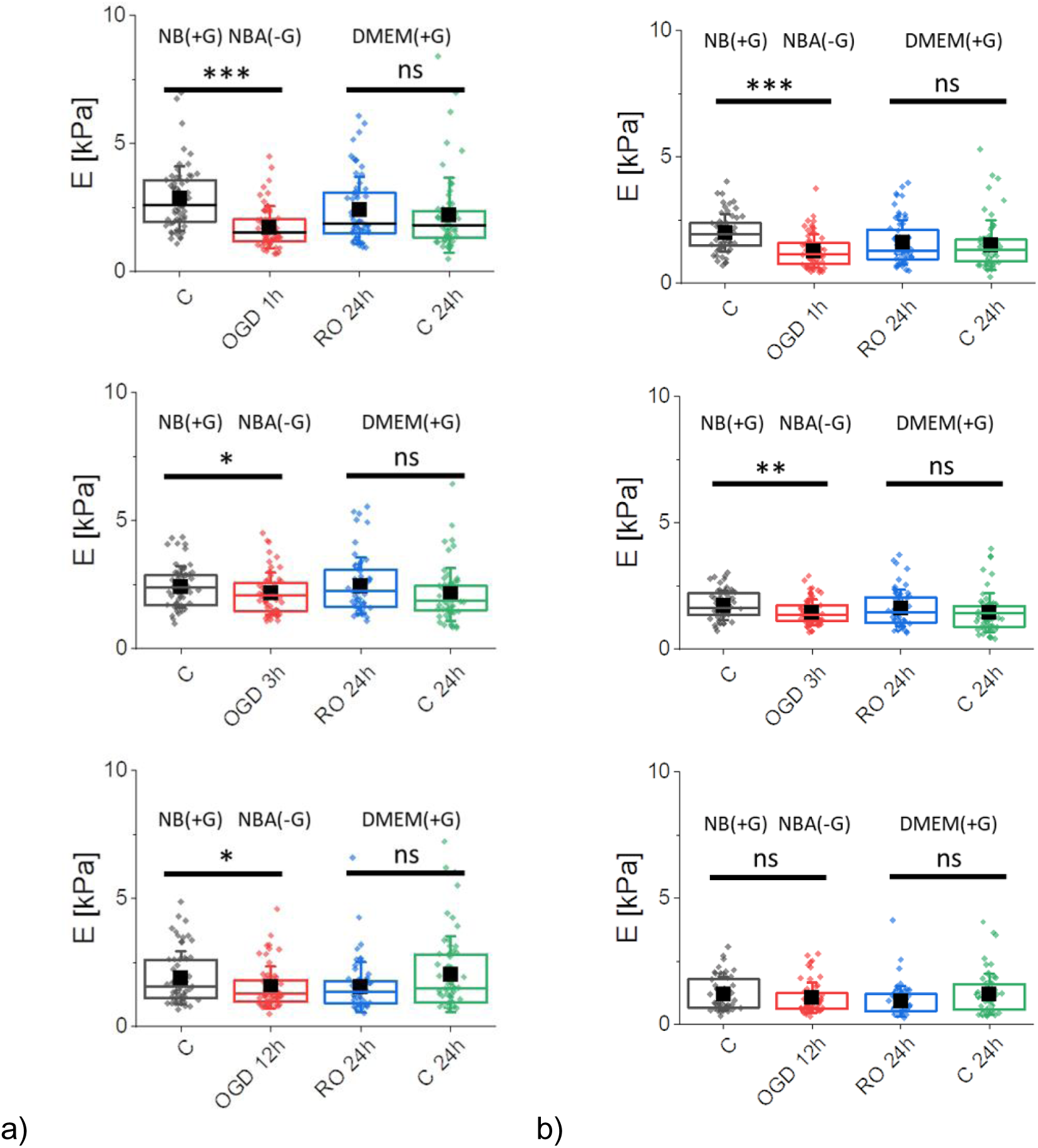
Nanomechanical properties of SH-SY5Y neuroblastoma cells after OGD treatment, quantified by the apparent Young’s modulus calculated for the indentation depth of 400 nm **(a)** and 1200 nm **(b)**. Four groups of cells were compared: control (C, NB(+G)), OGD cells (OGD 1h, 3h, or 12h, NBA(-G)), reoxygenated OGD cells (RO 24h, DMEM(+G)), and control, non-OGD cells (C 24h) kept in DMEM(+G) for the same time as reoxygenated OGD cells. Box plots represent a median (black line), a mean (solid square), standard deviation (whiskers), and 25% and 75% percentiles (box) from *n* = 60 cells. Statistical significance: ns – not statistically significant (p > 0.05), *p < 0.05, **p < 0.01, ***p < 0.001.

Young’s modulus was calculated at the shallow and deep indentations, i.e., 400 nm and 1200 nm, respectively. Alongside the already published data^22, 28, 35, 36^, the nanomechanical response of cells measured at shallow indentation (400 nm) reflects mainly the mechanics of the actin cortex. Thus, any alteration in cell mechanics can be related to the remodelling of actin filaments underlying beneath the cell membrane. AFM can measure only cells attached to the underlying surface; thus, to a certain extent, the mechanical properties of cells reflect the mechanics of cells resistant to unfavorable conditions. Cells heavily affected by OGD detached from the surface were not accessible for the AFM measurements. Still, in our study, mean values of Young’s modulus of OGD-treated cells significantly dropped by about 39.2% (*p* < 0.001), 10.7% (*p* = 0.045), and 19.4% (*p* = 0.042) for 1h, 3h, and 12h OGD in relation to control cells, respectively (**Fig. 2a**). Reoxygenation recovers the nanomechanical properties of cells close to values obtained for control, non-OGD cells kept in DMEM(+G) for 24h (same duration as reoxygenation). The elastic moduli are 2.40 ± 1.31 kPa versus 2.22 ± 1.46 kPa (*p* = 0.401), 2.46 ± 1.09 kPa versus 2.12 ± 1.03 kPa (*p* = 0.084), and 1.55 ± 0.98 kPa versus 1.05 ± 1.48 kPa (*p* = 0.110) for OGD-treated and non-treated cells, correspondingly. Thus, we can conclude that the recovery of the actin cortex occurs independently of the OGD duration. The largest changes were observed in cells after 1h OGD, but simultaneously, 24h of reoxygenation allowed cells to almost fully recover their mechanics. Longer OGD (3h and 12h) resulted in smaller mechanical changes than those observed after 1h of OGD.

The analysis of deeper indentations (like here 1200 nm) can evaluate the combined contributions of the actin cytoskeleton and other structural components of cells such as microtubules or cell nuclei. Mechanics of cells measured directly after OGD shows a significant drop after 1h and 3h. The apparent Young’s modulus dropped by 35.5% (*p* > 0.001) and 16.8% (*p* = 0.007), respectively (**Fig. 2b**). The OGD-induced mechanical changes were statistically insignificant after 12h of cell exposure to such conditions (*p* = 0.188). A similar level of changes suggests a weaker contribution of other cellular structures as compared to the actin cytoskeleton. In reoxygenated cells, changes in mechanical properties of OGD and non-OGD cells were statistically insignificant, showing a lack of mechanical contributions from deeper cellular layers. These results demonstrate that the mechanical response mainly contains the dominant contribution from the actin cytoskeleton.

### Organization of actin cytoskeleton in OGD-treated SH-SY5Y cells

Phase-contrast images collected prior to the AFM measurements did not show any particular changes in the macroscopic morphology. OGD treated cells reveal similar spindle and neuron-like morphology as control, non-treated cells, regardless of the OGD duration. As changes in cell mechanics are typically related to the organization of actin filaments, the confocal images with fluorescently labelled F-actin and cell nucleus were analyzed (**Fig. 4**).

**Figure 4.**
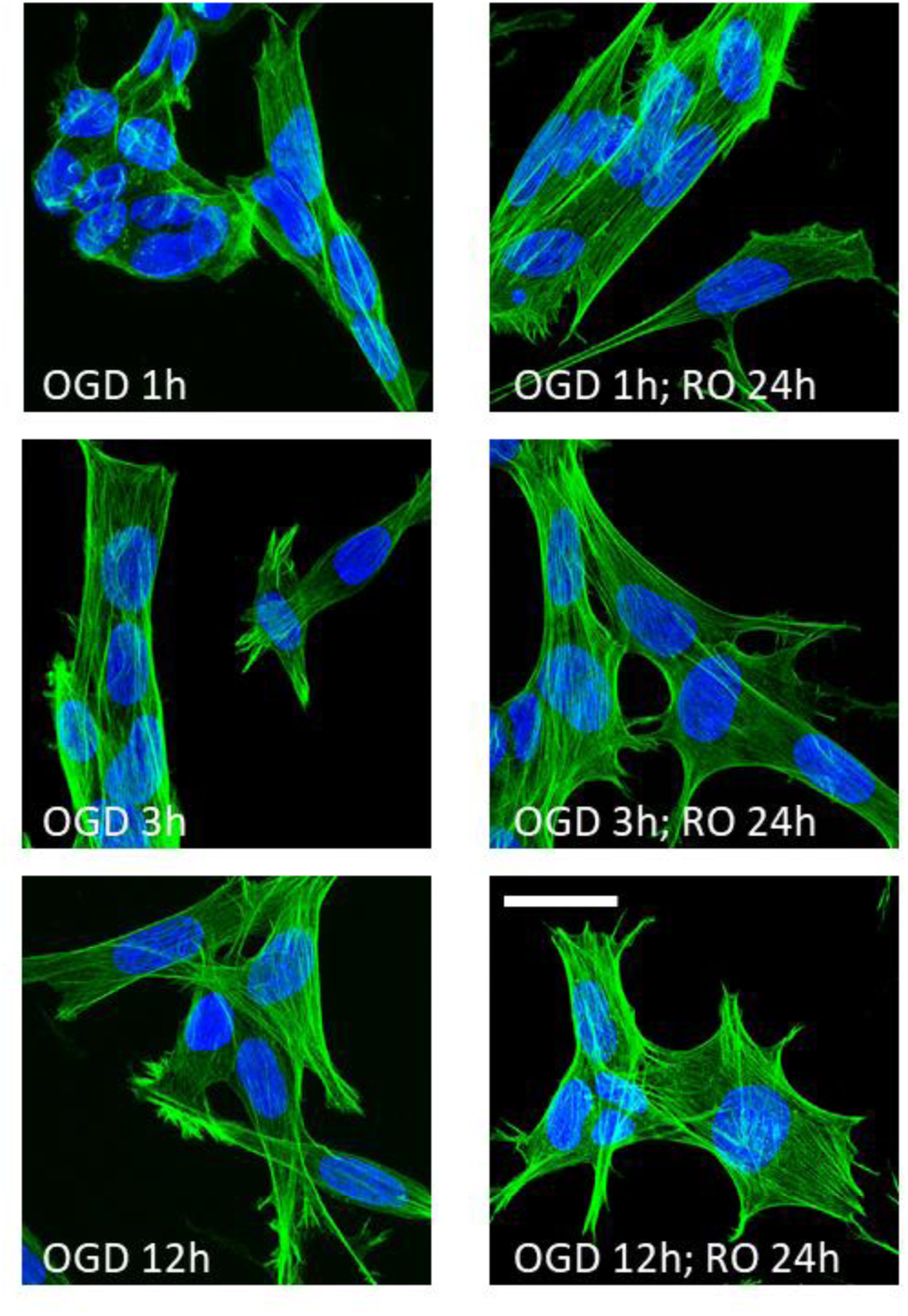
Confocal images of the actin cytoskeleton in OGD-treated and reoxygenated cells. Staining: actin filaments – phalloidin conjugated with Alexa Fluor 488, cell nuclei – Hoechst 33342; scale bar 25 µm.

In control and OGD treated cells, the organization of actin cytoskeleton was very similar, showing nicely actin bundles spanning over the whole cell. The only exception was cells visualized directly after 1h OGD, where cells change their morphology from a widely spread to a packed one (**Fig. 4**). This is consistent with the mechanical results showing the largest drop in the apparent Young’s (elastic) modulus. The organization of actin filaments in cells undergoing longer OGD treatment (3h and 12h) was barely visible, supporting weak changes in nanomechanical properties. Altogether, these results support the conclusion that the nanomechanical properties of cells are dominated by actin filament organization.

### Cell spreading as a measure of cell attachment

In our further steps, we performed a deeper analysis of the shape of individual cells using images recorded by epi-fluorescent microscopy (**Fig. 5**).

**Figure 5.**
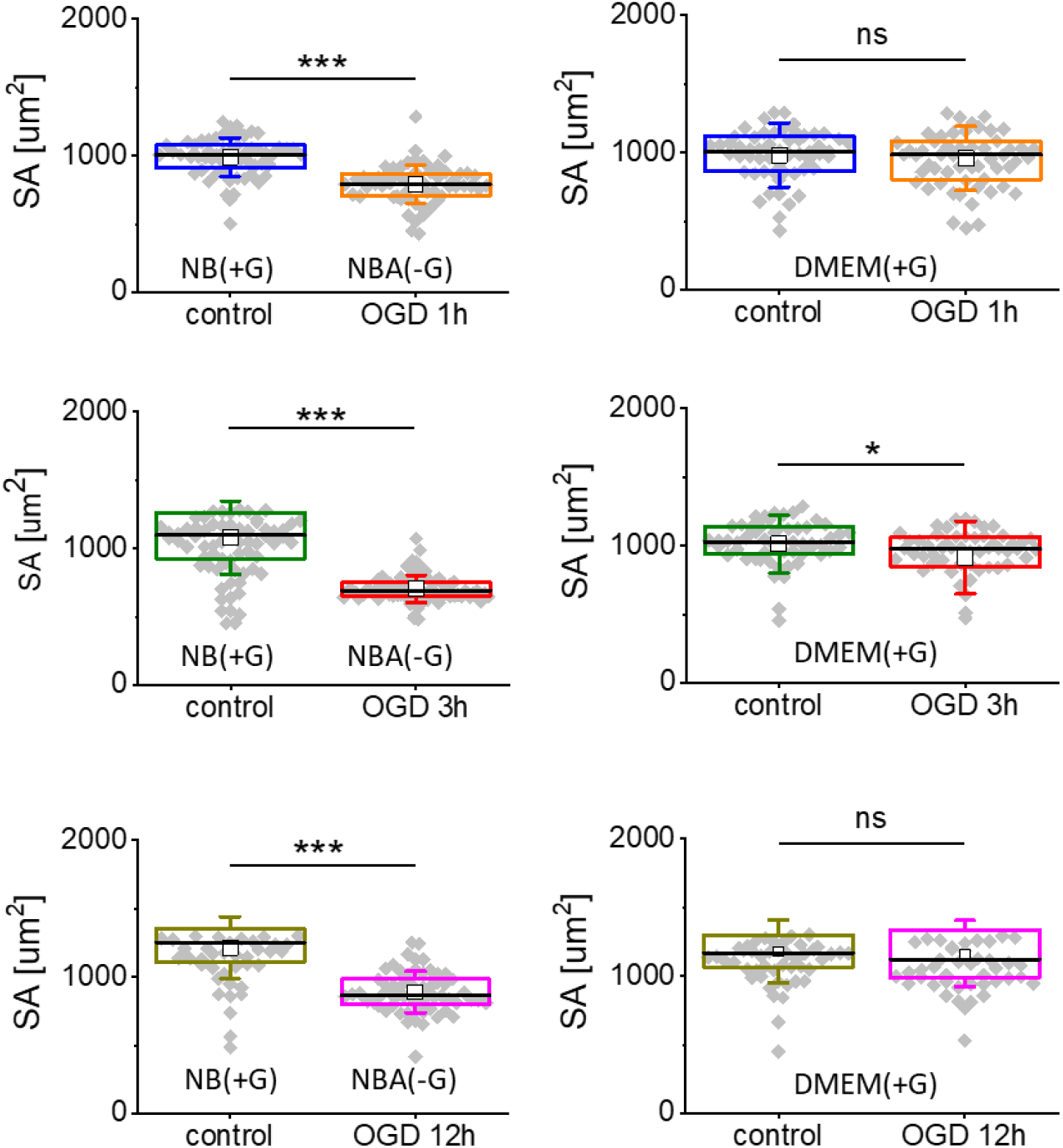
Spreading area (SA) of cells after OGD (a) and reoxygenation (b). Each dot denotes an average surface area of individual cells. Boxplot represents basic statistical parameters (mean, median, standard deviation, and 25% and 75% percentiles from n = 60 fluorescent images). Statistical significance: *ns* – not statistically significant (*p* > 0.05), *\*p* < 0.05, *\*\*\*p* < 0.001.

The results revealed that OGD-treated cells have different surface areas indicating impairments in their spreading on the surface (**Fig. 5a**). The smaller the surface area, the worse their attachments/adhesion to the surface is. The weak attachment of cells to the underlying surface was recovered after allowing cells to grow in reoxygenation conditions (**Fig. 5b**). The largest change in spreading area was observed for OGD-treated cells, while during reoxygenation, cells return to the surface area of control, non-treated cells. These results indicate that the spreading of cells involves the remodelling of actin filaments, which in our case is strongly related to the OGD treatment of SH-SY5Y cells.

### The shrinking of the cells is confirmed by the nucleus to cell ratio

Changes in cell surface area and lack of strong reorganization of the actin filaments suggest different mechanisms inducing alterations in nanomechanical properties of OGD-treated cells, such as changes in cell volume. A ratio between cell surface area (*C*) and cell nucleus (*N*) can quantify the latter (**Fig. 6**).

**Figure 6.**
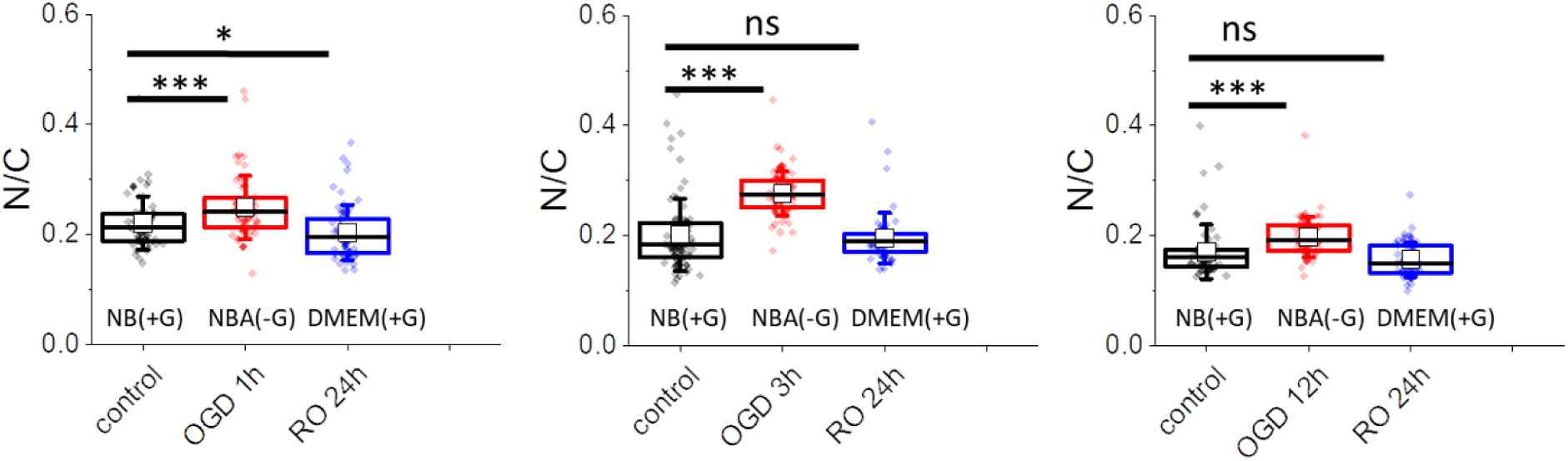
Nucleus to cytoplasm (*N/C*) ratio of cells after OGD treatment and reoxygenation. Each dot denotes an average value of individual cells. Boxplot represents basic statistical parameters, i.e., mean (open square), median (line), and standard deviation, from *n* = 60 cells. Statistical significance: *ns* – not statistically significant *\*p* < 0.05, *\*\*\*p* < 0.001.

The *N/C* value close to 1 indicates the dominant contribution of a cell nucleus in the surface area value, while its value close to 0 indicates a significant contribution from the cytoplasm.

The results show that the *N/C* ratio increases in cells visualized directly after OGD for all three tested timepoints and reaches the level of control cells after 24h of cell reoxygenation in all groups. Rought estimation of cell height from the cross-section of confocal images (from **Fig. 4**), shows that the height of the cell in the central area is 7.3 µm ± 1.4 µm (*n* = 14 cells), 8.3 µm ± 2.0 µm (*n* = 11) 10.9 µm ± 2.8 µm (*n* = 10) for cells after 1h, 3h, and 12h OGD, respectively. Altogether, these results show that the height of cells increases during OGD treatment.

### Cofilin level in OGD-treated SH-SY5Y cells

Cofilin is an actin regulating protein that quickly responds to various cell processes. Alterations in calcium ions, reactive oxygen species, ATP, or pH will result in quick dephosphorylation (activation) of cofilin^37^. Cofilin severs actin filaments but does not enhance actin depolymerization rate^38^. Instead, it creates new nucleation centres allowing for quick branching, polymerization, and depolymerization in a concentration-dependent manner^39^. The results of cofilin and p-cofilin expression levels in control and OGD treated cells are presented in **Fig. 7**.

**Figure 7.**
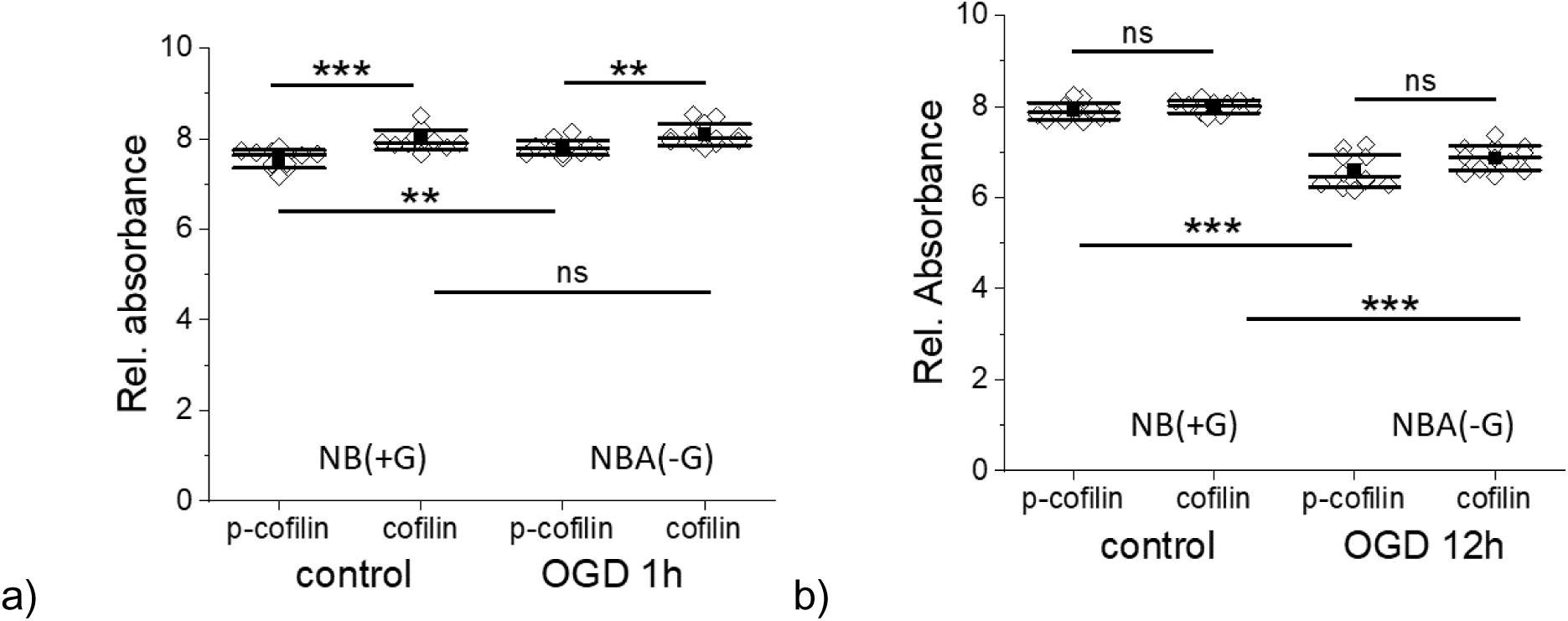
Cofilin and p-cofilin (phosphorylated cofilin) expression level in SH-SY5Y cells upon 1h **(a)** and 12h **(b)** OGD. Control cells were kept in NB(+G), while OGD cells were kept in NBA(-G). A mean (black square), median (black middle line), standard deviation (SD, outside black lines) were determined from data gathered from 3 independent repetitions. Statistical significance (*ns* – not statistically significant, *\*\*p* < 0.01, *\*\*\*p* < 0.001).

We observed the difference in the cofilin and p-cofilin levels in control cells that could be linked with a lower glucose level observed by proliferating cells in a given volume during a certain culture time^40^. When cofilin/p-cofilin was assessed in control cells simultaneously as cells after 1h of OGD, the concentration of cofilin was about 5% higher than p-cofilin (*p* < 0.001). In 1h OGD treated cells, the concentration of cofilin was only 3.5 % larger than p-cofilin (*p* < 0.002). In cells exposed to longer OGD duration, the expression level of cofilin in relation to p-cofilin vanishes (the same protein level was observed in cells after 3h and 12h OGD (**Fig. 7b** shows results of cofili/p-cofilin expression in SH-SY5Y cells after 12h OGD). Next, we compare the differences between the expression level of cofilins (or p-cofilins) for control and OGD treated cells. The determined *p*-values were *(i) p* = 0.006 (p-cofilin, between control and OGD (1h) cells) and *p* = 0.198 (cofilin, between control and OGD (1h) cells); *(ii) p <* 0.0001 (between control and OGD (3h) cells, regardless of the cofilin status); *(iii) p < 0.0001* (between control and OGD (3h) cells, regardless of the cofilin status). These results show that the ratio between cofilin/p-cofilin changes significantly during 1h OGD and vanish for a longer duration of OGD. Interestingly, the expression level of cofilin and p-cofilin in OGD cells decreases with OGD duration. Only, the expression level of p-cofilin changed.

## Discussion

Oxygen and glucose deprivation (OGD) is commonly used to study cerebral ischemic stroke. It mimics the process of a sudden disruption of blood flow to the brain. The lack of blood supply leads to decreased oxygen and glucose levels in the brain. The induced injuries activate various biochemical processes such as perturbation of calcium homeostasis^41^, malfunction of endoplasmic reticulum and mitochondria^42^, increased level of oxidative stress linked with DNA damage^43^. They directly affect cell morphology, which suggests changes in mechanical properties of OGD treated cells. In our work, AFM was applied to probe nanomechanical properties at the cellular level. Nanomechanics has already been reported to be altered during stroke^44^. The results have shown that tissue mechanics changes within the region affected by stroke and, also, at a distance from the stroke site^44^. AFM has shown the alterations in mechanical properties of the brain region severely affected by ischemia. Neuronal cells are mechanosensitive and highly responsive to altered mechanics of the surrounding environment ^45^. Thus, changes in the mechanical properties of ischemic tissue denote also changes in the functioning of neuronal cells.

Our study focused on the nanomechanical properties of SH-SY5Y cells subjected to OGD of different duration followed by 24-hour reoxygenation. AFM-based elasticity measurements were conducted at the shallow and deep indentations, which enabled us to quantify the changes occurring mainly in the network of actin filaments. A drop of Young’s modulus, a measure of cell deformability^22^, observed in cells subjected to OGD, suggests the reorganization of the cell cytoskeleton at the layer composed of the actin filaments. The most significant drop of Young’s modulus was observed in SH-SY5Y cells measured directly after OGD. When cells were allowed to grow in fully reoxygenated conditions, their elastic properties returned to the level of control cells. The cell metabolic activity (MTS assay) and cell viability (LDH assay) showed that the number of alive cells remained within 83-88% for both control and OGD treated cells. However, the metabolic activity of cells decreases with OGD duration, oppositely to changes observed in nanomechanical measurements.

Based on the obtained results, we propose the following mechanism leading to cell deformability changes during OGD. The observed time-dependent decrease of Young’s modulus in control, non-OGD treated SH-SY5Y cells was gradual regardless of the indentation depths chosen for the analysis (for low indentations of 400 nm and deeper indentations of 1200 nm). Probably, it reflects the impact of glucose consumption on the mechanical properties of cells. As the dynamic of assembly and disassembly of F-actin is strongly dependent on the accessibility of adenosine triphosphate (ATP) molecules (ATP-G-actin binds to barbed end three times faster than GTP-G-actin,^46^), the reduction of ATP resulted in slow disassembly of cytoskeleton resulting in a gradual decrease of Young’s modulus. A significantly lower level of ATP was reported to lead to cell softening^47^.

Several actin-associated proteins regulate actin polymerization/depolymerization as dynamic assembly and disassembly of the actin cytoskeleton is required for many biological processes, such as cell division, cell motility, endocytosis, and morphogenesis. These actin regulatory proteins contribute to nucleation, depolymerization, and fragmentation when a reorganization of the actin filaments is needed^48–51^. Cofilin is one of such proteins. It disassembles F-actin into tiny fragments of fibrous actin ^52^. Although severing of F-actin by cofilin is independent of energy addition, the extensive reorganization of actin filaments demands high energy supplies ^53^, in particular, to reach the level detected by AFM. It seems to be supported by the MTS assay showing unaltered metabolic activity of OGD-treated cells after one hour of the treatment. We postulate that in our measurements during the initial OGD, alterations in mechanical properties of cells reflect the disassembling effect of cofilin on the cortical actin. The effect is evident in cells after 1 hour of OGD. It is additionally detectable in the experiments, in which changes in the effective surface area of a single cell are quantified. The OGD duration of 1 hour is sufficient to induce a significant (∼20%) reduction of the mean single cell surface area and increase the cell height. Such increase suggested a more sparse actin scaffold and softening of the cells.

For longer OGD (3h and 12h), the cofilin-induced actin polymerization seems to be attenuated as changes in nanomechanical properties of OGD-treated cells are less pronounced. Severing and depolymerization of actin filaments by cofilin create many small F-actin fragments with new barbed ends needed for their polymerisation^54, 55^. Simultaneously, after prolonged OGD time, cells are metabolically impaired as the MTS assay reveals a significant drop in the number of metabolically active cells. It makes the reorganization of the actin cortex less favorable, indicating that cofilin activity is inhibited by phosphorylation of the serine residues at position 3 near the N-end for longer OGD duration. As a result, polymerization and stabilization of actin filaments are observed ^56^. Such effect leads to decreased cell deformability (cells become more rigid as Young’s modulus increases) in cells, in which metabolic activity is low. The latter affect the rebuilding of the actin filaments network, which is low, as shown by small values of the surface area of a single cell in cells after 3h and 12h of OGD.

After reoxygenation, the surface area went back to the levels of controls, but mechanical properties did not fully recover. When comparing short-time OGD, the elasticity of the cortical layer (400 nm) was well restored. The deep indentation, however, shows irreversible changes. Both shallow and deep indentations were irreversible after 12h OGD. The results contradict quantitative observation from actin and tubulin in fluorescence microscopy, where any significant change can be clearly observed. It indicates that changes in cell mechanics are more complex and cannot be explained only by actin (de) polymerization. Actin rearrangement via activation of profilin, cofilin and gelsolin, phosphorylation of myosin light chain, and changes in membrane spectrin cytoskeleton might be involved^57^. It should also be noted that in most studies, OGD constitutes the oxygen and glucose deprivations considered in parallel. Separating the specific oxygen- and glucose-related contributions calls for experiments conducted in conditions of either glucose or oxygen depletion^58^. The AFM-derived nanomechanical properties of OGD and non-OGD treated cells reveal a dominant role of glucose deprivation in 12 hours of OGD conditions that hinders the oxygen depletion effect.

Numerous research that use the OGD model to understand mechanisms involved in brain impairments^59, 60^ has demonstrated that the pathological process of ischemic stroke involves complex mechanisms acting on various cell types. These studies focus on cell or molecular biology aspects. Cofilin, being involved in the dynamic turnover of actin filaments, affects membrane integrity, receptor transport, and signal transduction. Understanding mechanisms responsible for cofilin-related changes of cytoskeleton remodeling, promise potential use of them to inhibit cofilin activity that might induce neuroprotection through targeting diverse cellular components and multiple pathways^61, 62^.

## Methods

### Cell culture

For experimental procedures, an undifferentiated SH-SY5Y human neuroblastoma cell line was used. Cells were cultured in a Dulbecco’s Modified Eagles’ Medium (DMEM, ATCC, LGC Standards) supplemented with 10% Fetal Bovine Serum (FBS, ATCC, LGC Standards). Cells were cultured in 35 cm^2^ culture flasks (TPP) and passaged (< 10) into the corresponding plastic media required in each experiment. Cells were culture in the CO_2_ incubator (NUAIRE) at 37°C and 5%CO_2_/95% air atmosphere.

### OGD experiments

Cells were passaged from the culture flask to the Petri dish (TPP) and kept in the CO_2_ (37°C and 5%CO_2_/95% air atmosphere) for 24 hours in the DMEM (ATCC, LGC Standards) supplemented with 10% FBS. DMEM contains 4500 mg/L glucose and 1 mM sodium pyruvate. After this time, the medium was replaced either with (1) neurobasal medium (NB) containing glucose (control cells, undergoing the same treatment as OGD cells but without applying OGD conditions) or with neurobasal A medium (NBA, without glucose used to create OGD conditions). NB is optimized for prenatal and fetal neurons. NBA is optimized for growing postnatal and adult brain neurons. These two media differ only in osmolality (260 mOsm versus 235 mOsm, respectively).

OGD conditions were obtained in the following way. Cells were placed in the temperature-controlled table CO_2_ incubator (Olympus) at 37°C. The incubator was connected to a gas exchange 3-input system (Tokai Hit) supplying air, N_2_, and CO_2_. In our system, CO_2_ concentration remained constant (5%) while the air was replaced by N_2_, resulting in an oxygen concentration of 0.1%. The oxygen level was maintained constant by applying a gas flow at a level of 150 ml/min. These parameters were kept constant for 1, 3, and 12 hours. Immediately after applying OGD, cells were analyzed using various techniques: MTS and LDH assays, atomic force microscopy (AFM), epifluorescence, and confocal microscopy.

### MTS assay

The viability and metabolic activity of SH-5YSY cells were verified by using an MTS colorimetric test (Promega). Cells were cultured in 24-well plates in 1 ml of the culture medium (DMEM). Next, 100 µL of MTS reagent (tetrazolium compound) was added to the cells. Then, cells were incubated at 37◦C in 95% air/5% CO2 atmosphere in the CO2 incubator (Nuaire) for 2 h. The MTS method reduces tetrazolium compounds by viable cells to generate a colored formazan product soluble in cell culture media. The final volume of 1.1 mL was pipetted to 96-well plates with 100 µL per hole. The absorbance (OD = 490 nm) was recorded for 0h, 3h, and 24h after OGD using a spectrophotometer (ELISA SPECTROstar Nano, BMG LABTECH). The MTS assay was repeated three times.

### LDH assay

The cytotoxic effect of OGD was quantified by using CyQUANT™ LDH Cytotoxicity Assay Kit (Invitrogen). Cells were plated on 24-well plates in 1 ml of the corresponding culture medium (**Fig. 1**). LDH level was evaluated in the following samples, i.e., (i) control and OGD cells, (ii) control and OGD cells treated with lysis buffer (10% by volume, 45 min in the CO_2_ incubator), (ii) culture medium (supernatant) taken from control or OGD cells and separately treated with lysis buffer in an analogous way as (ii) to verify how much cells detached during the medium exchange. Then, 50 µl of the medium from each sample type was aspirated from each well and transferred into a 96-well plate. Next, to each well, 50 µl of the reaction mixture was added. After 1h of incubation in conditions protecting against light exposure, 50 µl of stop solution was added. Lactate dehydrogenase (LDH) is a cytosolic enzyme present in various cell types. Damage of cell membrane results in a release of LDH to the surrounding medium, which can be quantified by using LDH as a catalytic enzyme. It converts lactate to pyruvate *via* NAD+ reduction to NADH. Oxidation of NADH by diphosphorase reduces a tetrazolium salt to a red formazan product. OD at 490nm was registered using an ELISA reader (Ledetect 96 ELISA, LED-based microplate reader, Labexim Products) to detect it. The level of formazan is directly proportional to the level of LDH in the surrounding medium. To obtain cell viability level the following equation was used:

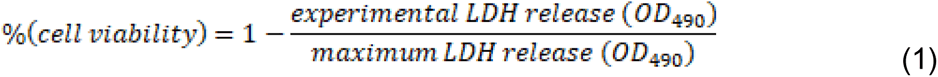

where the maximum LDH release is the sum of LDH release in Triton X100 treated samples (cells and supernatants of control and OGD-treated cells, respectively).

### Phospho-Cofilin/Cofilin assay

To obtain changes in cofilin activity level, CytoGlow™ Cofilin (Phospho-Ser3) Colorimetric Cell-Based ELISA Kit was applied (Assay Biotechnology) to monitor target proteins concentration, here, in cells undergoing OGD treatment. Briefly, SH-SY5Y cells (50,000 per well) were plated on a 96-well plate. After OGD experiments, cells were fixed using 4% paraformaldehyde and washed three times with 200 µl with Wash Buffer (WB) for 5 minutes, each time gentle shook. Then, 100 µl of quenching solution was added for 20 minutes at room temperature (RT), followed by 3 times washing with WB for 5 minutes at a time. Next, 200 µl of Blocking Buffer was added for 1 hour at RT, and, afterwards, the plate was washed again (3 x times, WB at RT). Then, a solution of 50 µl of each primary antibody against phosphorylated cofilin (Anti-Cofilin (Phospho-Ser3) antibody), cofilin (Anti-cofilin antibody) and Glyceraldehyde 3-phosphate dehydrogenase, GADPH (Anti-GAPDH antibody) was added to the corresponding well and incubated for 16 hours (overnight) at 4°C. Afterwards, they were rinsed 3 times with 200 µl of WB for 5 minutes. In the next step, secondary antibodies (horseradish peroxidase (HRP)-conjugated antiRabbit IgG antibody and/or HRP-conjugated anti-Mouse IgG antibody) were added (50 µl) for 1.5 incubation at RT. After incubation, the plate with cells was washed, and 50 µl of Ready-to-Use Substrate was added to each well for 30 minutes at RT, followed by adding a Stop Solution. OD at 450 nm was immediately read using a microplate reader (ELISA SPECTROstar Nano, BMG LABTECH).

### Atomic force microscope (AFM)

The mechanical properties of cells were measured using AFM (CellHesion, Bruker-JPK, Germany). The microscope is equipped with a constant temperature system. In our experiments, the temperature was set to 32°C to provide the cell survival conditions and the cantilever stability. Cells were indented with silicon nitride cantilevers (ORC-8, Bruker) characterized by a nominal spring constant of 0.03 N/m and an open half-angle of 36°. All measurements were conducted in a force spectroscopy mode. The spring constants of used cantilevers were determined using the Sader method ^63^. The average value was 0.058 ± 0.005 for *n* = 8 cantilevers. A force map of 6 per 6 pixels (corresponding to a 6 µm x 6 µm scan size) was recorded on each cell.

The force curves (i.e. the dependence of the cantilever deflection recorded as a function of relative sample position) were acquired at the approach/retract velocity of 8 µm/s. On individual plastic Petri dishes, two groups of force curves were collected. First, calibration curves were acquired on a Petri dish bottom surface (a reference calibration curve). Next, force curves were recorded on living cells. Next, force curves were recorded on living cells. Force curves were collected by setting a grid of 6 × 6 points that corresponded to 6 µm × 6 µm scan area. A grid was set over the nuclear region to minimize the influence of the underlying stiff plastic surface. All measurements were conducted in DMEM and were repeated 3 times.

### Young’s modulus determination

The subtraction of the calibration curves from a curve collected on a living cell produces the relation between load force and indentations depth. This relation was analyzed using the Hertz-Sneddon contact mechanics. The AFM probe was approximated by the cone that resulted in the following relation between load force and indentation depth:

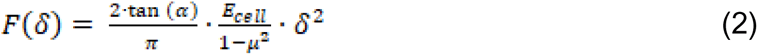

where *F* is load force, *δ* is the indentation depth, Ecell is the apparent Young’s modulus of the cell, and *µ* is the Poisson’s ratio (equalled to 0.5 assuming that cells are incompressible materials). The final modulus value was expressed as a mean and standard deviation from all measured cells.

### Fluorescence (epifluorescence and confocal) microscopy

Visualization of actin filaments, microtubules, and cell nuclei was performed by using Olympus IX83 (Olympus, Japan) fluorescence microscope equipped in objectives 20x and 40x magnification, 100 W mercury lamp (illuminating the whole cell area uniformly), and a set of filters to record emission at 594 nm and 420 nm. Images were collected using Orca Spark digital camera providing a 2.3 megapixel (1920x1200) pixel image and analyzed with ImageJ (ImageJ 1.53e https://imagej.nih.gov/ij/). Cells cultured on 24-well plates were fixed in 3.7% paraformaldehyde, then they were washed with phosphate-buffered saline (PBS, Sigma), treated with a cold 0.2% Triton X-100 solution, and again washed with the PBS buffer. Afterwards, cells were incubated with β-tubulin antibody conjugated with Cy3 for 24 hours. The next day, samples were stained with phalloidin conjugated with AlexaFluor 488 dye during 1h incubation. Cell nuclei were stained by 10 min incubation with Hoechst dye.

Confocal images of actin and microtubular cytoskeleton were recorded at the Laboratory of in vivo and in vitro Imaging (Maj Institute of Pharmacology Polish Academy of Science, Cracow, Poland). They were recorded using a Leica TCS SP8 WLL confocal microscope equipped with new-generation HyD detectors set at 415-450 nm (Hoechst) and 509-560 nm (Alexa Fluor 488). Fluorescent dyes were excited by diode lasers: 405 nm (Hoechst) and white light laser with emission wavelength set at 499 nm (AlexaFluor 488). Images were registered using an oil immersion 63x objective lens (HC PL APO CS2 NA 1.40).

### Surface area determination

A single cell effective surface area (SA) was applied to characterize how well cells spread on the surface at given conditions. This value describes an average surface area occupied by an individual cell. Images of fluorescently stained cells (F-actin using phalloidin-Alexa Fluor 488 dye, cell nuclei by Hoechst 33342) were binarised using ImageJ software. From these images, the surface area occupied by cells was determined. Next, cell nuclei were manually counted to receive the number of cells. Finally, the surface area occupied by cells was divided by the number of cells that enabled the calculation of the effective surface area of a single cell. Images were acquired during three repetitive experiments, which resulted in 20 images per condition to be analyzed. The total number of cells was at least 8000 cells.

### Nucleus – to – cytoplasm (N/C ratio)

To obtain the N/C, the effective area of individual cell nuclei was quantified analogously as the effective surface area of a single cell was determined. Next, the effective surface area of a single nucleus was divided by the effective surface area of a single cell. The total number of images analyzed was 20 per condition.

### Statistical analysis

All data are presented as the mean ± standard deviation from n repetitions. In all figures, box plots were applied to show the basic statistical descriptors: mean (open square), median (black line), standard deviation (whiskers), and 25% and 75% percentiles (box). Statistical significance was verified by applying the non-parametric Mann-Whitney test (Origin 9.2 Pro). Significance is indicated by p values (ns – not statistically different, p > 0.05; *p < 0.05, **p < 0.01, ***p < 0.001).

## Acknowledgments

TZ acknowledges the support of project no. POWR.03.02.00-00-I013/16 (InterDokMed). The APC was funded by project no. POWR.03.02.00-00-I013/16 (InterDokMed)". The authors are thankful to prof. Danuta Jantas and prof. Halina Jurkowska for sharing the SH-SY5Y cell lines.

## Author Contributions

Conceptualization, TZ, ML, JP and BZ; methodology, TZ and ML; validation, TZ, BZ, JP and ML; formal analysis, TZ; investigation, TZ; resources, ML; data curation, TZ; writing - original draft preparation, TZ, BZ, JP, ML; writing—review and editing TZ, BZ, JP, ML; confocal images visualization, JW; cell cultures, JP; supervision, ML and JP; funding acquisition, ML; All authors have read and agreed to the published version of the manuscript.

## Data Availability Statement

**Correspondence** and requests for materials should be addressed to ML.

